# The genomic and proteomic landscape of the rumen microbiome revealed by comprehensive genome-resolved metagenomics

**DOI:** 10.1101/489443

**Authors:** Robert D. Stewart, Marc D. Auffret, Amanda Warr, Alan W. Walker, Rainer Roehe, Mick Watson

## Abstract

Ruminants provide essential nutrition for billions of people worldwide. The rumen is a specialised stomach adapted to the breakdown of plant-derived complex polysaccharides, and collectively the rumen microbiota encode the thousands of enzymes responsible. Here we present a comprehensive analysis of over 6.5 terabytes of Illumina and Nanopore sequence data, including assembly of 4941 metagenome-assembled genomes, and several single-contig, whole-chromosome assemblies of novel rumen bacteria. We also present the largest dataset of predicted proteins from the rumen, and provide rich annotation against public datasets. Together these data will form an essential part of future studies of rumen microbiome structure and function.

## Introduction

Ruminants are one of the most important food-producing species on the planet, providing food to billions of people worldwide. Crucially, ruminants convert human-inedible, low value plant biomass into products of high nutritional value, such as meat and dairy products. Ruminants extract energy and nutrients from recalcitrant plant material via microbial fermentation in the rumen, the first of four chambers of the stomach. Here, a mixture of bacteria, archaea, fungi and protozoa ferment complex carbohydrates (e.g. lignocellulose, cellulose) to produce short-chain fatty acids (SCFAs) that the host animal uses for homeostasis and growth. As such, ruminant microbes are a fertile hunting ground for novel enzymes efficient at plant biomass degradation for use in the biofuels industry^1–3^, and manipulation of the rumen microbiome offers opportunities for lowering the cost of food production^4^.

The success of ruminants is of importance to two major issues facing humanity. The first is food security – the world population is growing, driven largely by lower- and middle-income countries (LMICs), and we are under pressure to produce more food whilst using fewer resources. This is a daunting challenge, and the agricultural system must respond by increasing the efficiency of food production. The second is climate change – an estimated 14% of methane produced by human activity has been attributed to ruminant livestock, released annually into the atmosphere^5^. Methane is a natural by-product of ruminant fermentation, released by methanogenic archaea. This production represents a loss of energy and carbon for the animal, estimated to be between 2 and 12%^6^. Methane production has been directly linked to the abundance of methanogenic archaea in the rumen^7^,^8^, and is also under the control of rumen microbial metabolism thermodynamically dependent of hydrogen partial pressure^9^. Moreover, it is been shown to be under the influence of host genetics^10^, offering possibilities for mitigating this issue through selection or manipulation of the microbiome.

Recently, two studies published the genomes of over 1300 rumen microbes. Firstly, we released 913 draft and near-complete metagenome-assembled genomes (MAGs) from the rumen of 43 cattle raised in Scotland^11^. We coined the phrase “rumen-uncultured genomes”, or RUGs, to refer to these. Using these genomes we showed that classification rates for rumen metagenomic sequencing data could be increased from 7% to an average of 50%; that there remain many novel rumen microbial species; and that the genomes of rumen microbes harbour millions of novel proteins, many involved in carbohydrate metabolism. Soon after, 410 genomes from the Hungate collection were sequenced and released^12^, and described alongside 93 additional rumen microbial reference genomes in the public domain. As isolate-genomes, the Hungate collection are more complete, and crucially, exist in culture, so can be grown and studied in the lab. However, we found that the addition of the Hungate collection genomes only increased metagenomic classification rates by 2-4 percent, suggesting that there are many more un-cultured and un-sequenced microbes in the rumen of modern cattle breeds.

We are in the midst of a metagenomic revolution, driven by affordable, high-quality sequence data^13^,^14^, and advances in bioinformatics and computational power^15–19^. Despite the fragmented assemblies produced, even short-read Illumina data can be efficiently and accurately binned into high quality genomes. As a result, many previously under-studied microbial environments now have an abundance of MAGs – examples include 913 MAGs from the rumen^11^, 957 from the ocean^20^, 2540 from underground aquifers^21^, and almost 8000 from a range of environments covered by public sequence data^22^. Although these numbers are not directly comparable due to the use of different quality metrics, the scale of the datasets emphasizes the power of metagenome-resolved metagenomics. Recent attention has focused on the use of single-molecule, long-read sequencing technologies, such as PacBio^23^ and Oxford Nanopore^24^, for metagenomic analysis ^25^. It is known that such technologies can accurately and efficiently assemble whole bacterial chromosomes^26–29^, but their application to complex metagenomes has not yet been demonstrated.

In this paper we present comprehensive metagenomic analysis of over 6.5Tb of sequence data from the rumen of 282 cattle. Our genome-resolved metagenomics workflow produced 4941 de-replicated, draft and near-complete bacterial and archaeal genomes, including 4056 genomes not present in our previous dataset, and 312 updated genomes of higher quality. We also present a metagenomic assembly of Nanopore sequencing data from one of the samples. This assembly contains at least three whole bacterial chromosomes as single contigs and represents the most continuous metagenomic assembly from the rumen to date.

We demonstrate that our new RUG dataset contains thousands of novel microbial genomes that encode millions of novel proteins differing significantly from previously published rumen genomic and metagenomic datasets. The genomic and proteomic resources presented here will form an important reference dataset for future studies on the structure and function of the rumen microbiome.

## Results

### 4941 metagenome-assembled genomes from the cow rumen

We have sequenced DNA extracted from the rumen contents of 282 beef cattle, producing over 6.5Tb of Illumina sequence data. Here we present an analysis of 4941 rumen uncultured genomes (RUGs) including 4056 new genomes that did not appear in our previous dataset, and 312 updated genomes of higher quality. We operate a continuous assembly-and-de-replication pipeline, which means that newer genomes of the same strain (>99% average-nucleotide identity, ANI) can replace older genomes if their completeness and contamination statistics are better. All of the 4941 RUGs we report here have completeness >= 80% and contamination <= 10%.

The 4941 RUGs were analysed using MAGpy^19^ – assembly characteristics, putative names and taxonomic classifications are in Supplementary data 1, and Sourmash^30^, DIAMOND^31^ and PhyloPhlAn^32^ results are in Supplementary data 2. A phylogenetic tree of the 4941 RUGs alongside 460 public genomes from the Hungate 1000 project can be seen in Figure 1 and Supplementary data 3. The tree is dominated by large numbers of genomes from the *Firmicutes* and *Bacteroidetes* phyla (dominated by *Clostridiales* and *Bacteroidales* respectively), but also contains many novel genomes from the *Actinobacteria, Fibrobacteres* and *Proteobacteria* phyla. *Clostridiales* (2079) and *Bacteroidales* (1081) are the dominant orders, with *Ruminoccocacae* (1111) and *Lachnospiraceae* (640) the dominant families within *Clostridiales* and *Prevotellaceae* (521) the dominant family within the *Bacteroidales*.

**Figure 1.**
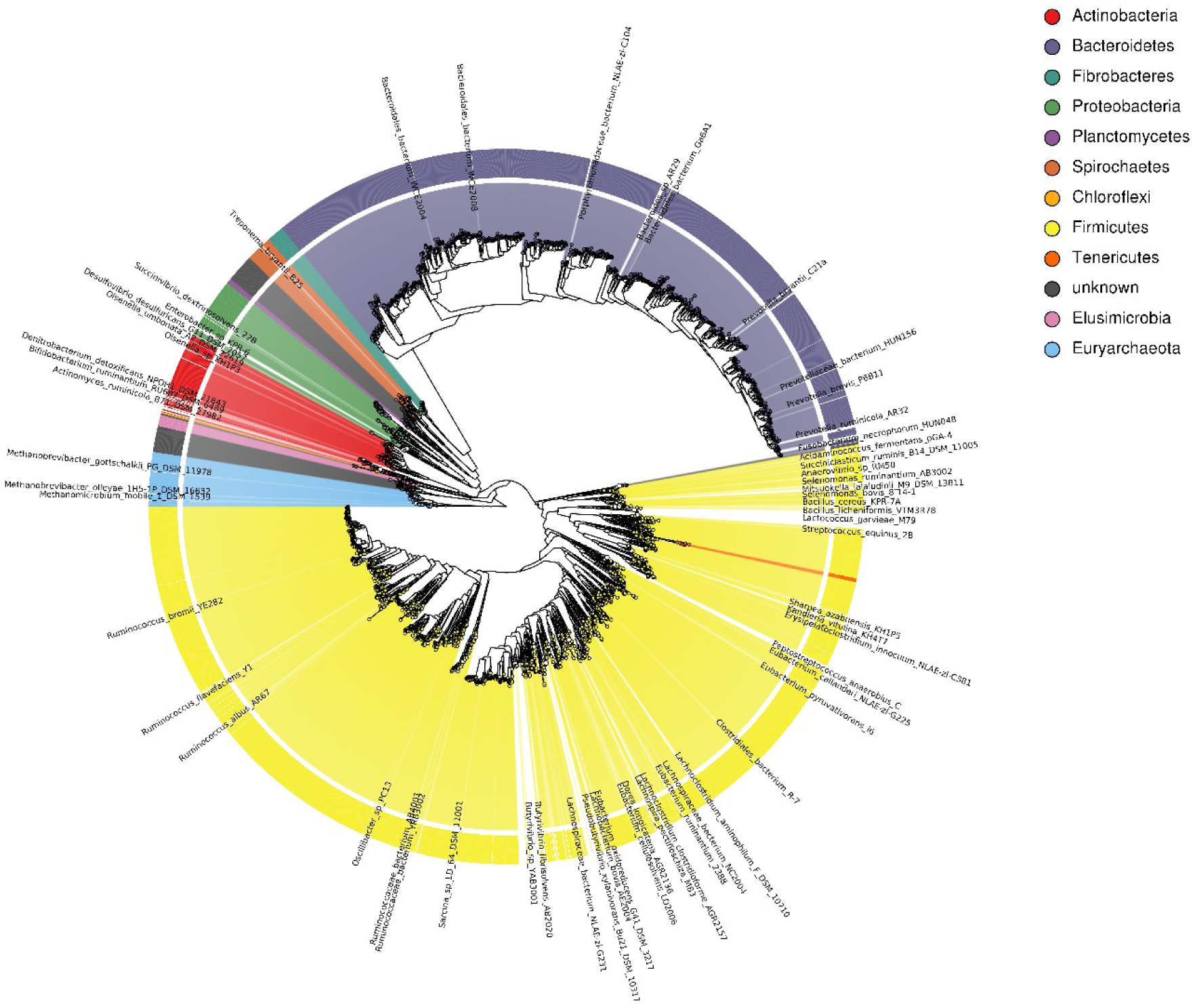
Phylogenetic tree of 4941 rumen uncultured genomes (RUGs) from the cow rumen, also incorporating 460 rumen genomes from the Hungate 1000 collection. Tree produced by PhyloPhlAn^32^.

Of the 4941 genomes, 144 are classified to species level, 1092 to genus level, 3188 to family, 4084 to order, 4514 to class, 4801 to phylum and 4941 to kingdom. Of the genomes classified at the species level, 43 represent genomes derived from uncultured strains of *Ruminococcus flavefaciens*, 42 represent genomes from uncultured strains of *Fibrobacter succinogenes*, 18 represent genomes from uncultured strains of *Sharpea azabuensis*, and 10 represent genomes from uncultured strains of *Selenomonas ruminantium*. These species belong to genera known to play an important role in rumen homeostasis^33^.

There are 126 Archaeal genomes, 111 of which appear to be species of *Methanobrevibacter*. However, 4 of the RUGs match more closely to the unclassified *Thermoplasmata* “methanogenic archaeon iso4-h5” known to be a ruminal representative of the methylotrophic methanogens^34^, whereas nine others form a separate group, and are listed as “uncultured euryarchaeota”. These nine therefore represent previously undiscovered ruminant archaea.

Genome quality statistics can be seen in Figure 2. There are competing standards for the definition of MAG quality - for example, Bowers *et al* ^35^ describe high-quality drafts as having >= 90% completeness and <= 5% contamination; 2417 of the RUGs meet these criteria. Alternatively, Parks *et al* ^22^ define a quality score as “Completeness – (5 * contamination)” and exclude any MAG with a score less than 50; 4761 of the RUGs meet that criterion; however, whilst the MAGs from Parks *et al* could be as low as 50% complete, the genomes presented here are all >= 80% complete.

**Figure 2.**
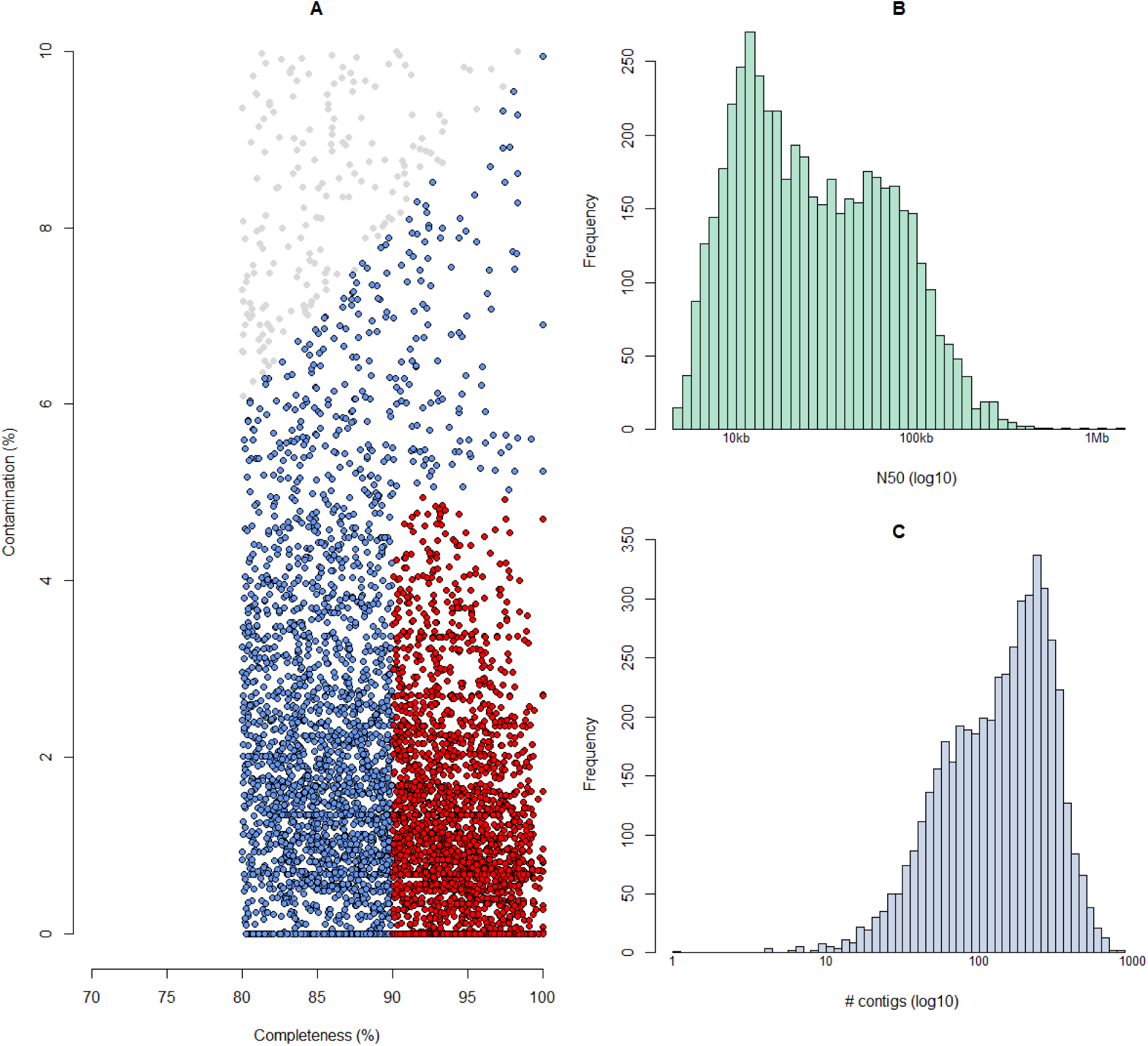
Assembly quality. **A)** Completeness and contamination statistics for 4941 RUGs. Red points indicate the highest quality genomes with >=90% completeness and <=5% contamination. All other RUGs are >80% complete and <=10% contaminated. Those in blue have a quality score >=50 as defined by Parks *et al*, whereas those in grey have a quality score <= 50. **B)** Histogram of N50 for 4941 RUGs (log10 scale). **C)** Histogram of the number of contigs per genome for 4941 RUGs.

The RUGs range in size from 456kb to 6.6Mb with N50s (50% of assembled bases are in contigs greater than the N50 value) ranging from 4.5kb to 1.37Mb. Metagenomic assembly of Illumina data often results in highly fragmented assemblies – however, surprisingly, one of our RUGs is a single contig of just over 1Mb in size. RUG14498 is an uncultured *Proteobacteria* with an estimated completeness value of 87.91% and contamination value of 0% (136 of 147 single-copy genes present with no duplications). *Proteobacteria* with small genomes (<1.5Mb size) are relatively common (n=67) in our dataset and have also been found in other large meta-genome assembly projects^22^. The *Proteobacteria* genomes we present here are undoubtedly novel, showing only between 45% and 60% amino acid identity with proteins in UniProt TREMBL^36^. We compared our single-contig *Proteobacteria* assembly with nine *Proteobacteria* with similarly sized genomes assembled by Parks *et al* ^22^, and whole-genome alignments can be seen in Figure 3. There are clear, linear, whole-genome alignments between RUG14498 (on the x-axis) and several of the UBA (“Uncultivated Bacteria and Archaea”) MAGs, suggesting that the single-contig RUG14498 is a high-quality, near-complete whole genome of a novel *Proteobacteria* species. Two other long Illumina contigs exist, 1.37Mb and 1.38Mb in size respectively, both from multi-contig unknown *Bacteroidia* RUG assemblies.

**Figure 3.**
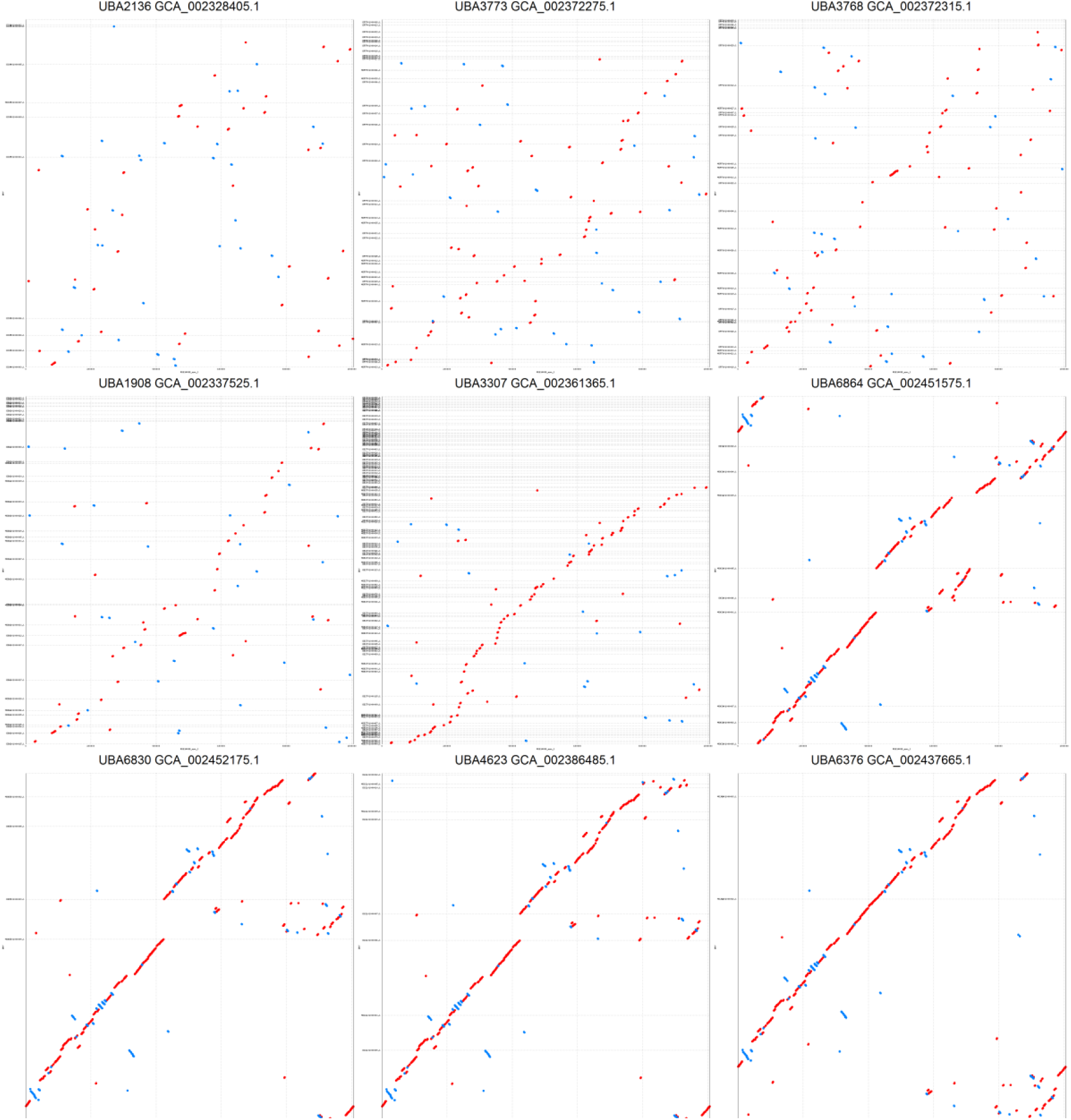
Whole genome alignments between the single-contig Illumina assembly RUG14498 (the x-axis on all plots) and nine similarly sized *Proteobacteria* MAGs from Parks *et al*. Clear, linear whole-genome alignments between RUG14498 and six of the Parks *et al*^22^ MAGs can be seen, with faint linear alignments distinguishable on a further two. UBA3307 and UBA1908 appear to include additional sequence with no orthologous matches in RUG14498.

RUG14498 has a single full length 16S rRNA gene (1507bp). The top hit in GenBank (97% identity across 99% of the length) is accession AB824499.1, a sequence from an “uncultured bacterium” from “the rumen of Thai native cattle and swamp buffaloes”. The top hit in SILVA^37^ is to the same sequence, only this time annotated as an uncultured *Rhodospirillales*. The average protein identity between RUG14498 and UniProt TREMBL is 45.59%, and this together with the 16S rRNA gene results above supports the conclusion that RUG14498 represents a novel *Proteobacteria* species within the *Rhodospirillales* order. Such low amino acid identity to known proteins limits our ability to predict function and metabolic activity; nevertheless RUG14498 appears to encode 73 CAZymes, including 42 glycosyl transferases and 19 glycosyl hydrolases, suggesting a role in carbohydrate synthesis and metabolism.

There are further near-complete genomes in the RUGs. In addition to the single-contig RUG14498, 22 other genomes have <= 10 contigs, and 98 have <=20 contigs. Unfortunately, many of these do not have close matches in either GenBank or the Hungate 1000 collection, limiting our ability to carry out comparative genomic analysis. However, RUG13721 has only 25 contigs, is estimated 99.77% complete and 0.86% contaminated, and shows high protein identity (95.94%) to *Bifidobacterium merycicum* DSM6492, part of the Hungate collection. *Bifidobacterium merycicum* DSM6492 has only slightly better assembly characteristics than RUG13721 (15 contigs vs 25 for RUG13721; 647kb N50 vs 156kb for RUG13721), and whole-genome alignments show that RUG13721 accurately captures the full genome of *Bifidobacterium merycicum* (Figure 4).

**Figure 4.**
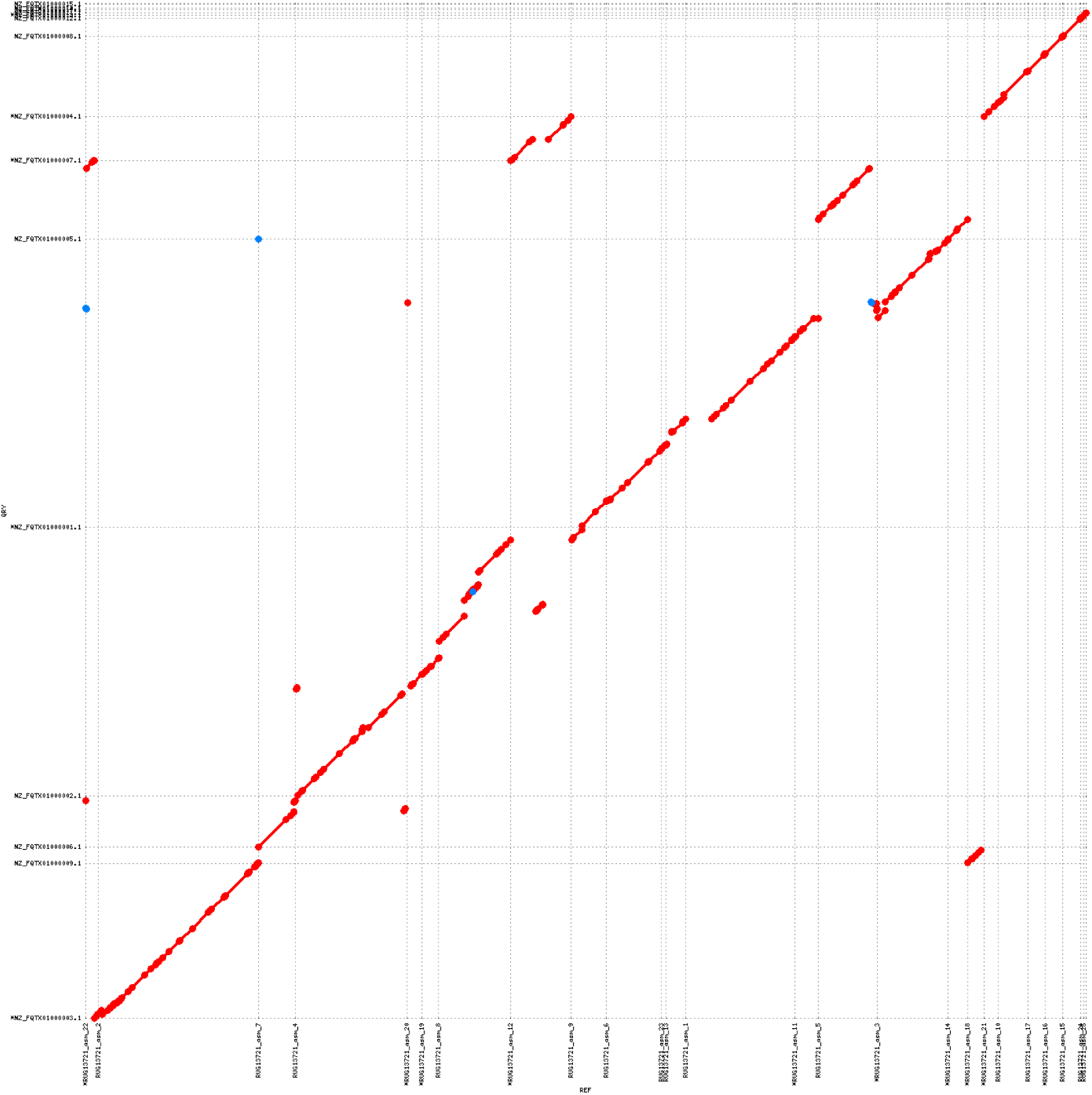
Whole-genome alignment between RUG13721 and *Bifidobacterium merycicum* DSM6492. Both genomes exist as unordered contigs. MUMmer has made an attempt to re-order the contigs to find the best match

To demonstrate the novelty of the 4941 RUGs, we compared them to both the Hungate collection and our previous dataset^11^ – the results are in Figure 5. Overall, the RUGS represent significant novelty compared to the Hungate collection (Figure 5A). Only 149 of the RUGs share greater than 95% protein identity with Hungate members; only 271 show greater than 90%; and therefore 4670 of the RUGs show less than 90% protein identity with Hungate member strains. As might be expected, there are higher levels of similarity shown between the 4941 RUG dataset presented in this paper, and our smaller dataset previously published (Figure 5B). However, still 2387 of the RUGs show less than 90% protein identity, and over 1100 of the RUGs show less than 70% protein identity, compared with Stewart et al. There is a clear inflection point in Figure 5B, roughly half way along the x-axis, where the protein identity dips below 90% and the Mash distance rises, neatly demonstrating the novelty represented by our new, larger dataset. Many of the RUGs at the lower end of the scale could not be classified beyond Phylum level, and some are simply “uncultured bacterium”. Taken together, these statistics demonstrate that there is significant novelty in the larger 4941 RUG dataset as compared to previously published rumen genomes, and that there remain thousands of novel species of bacteria and archaea within the rumen that have yet to be cultured.

**Figure 5.**
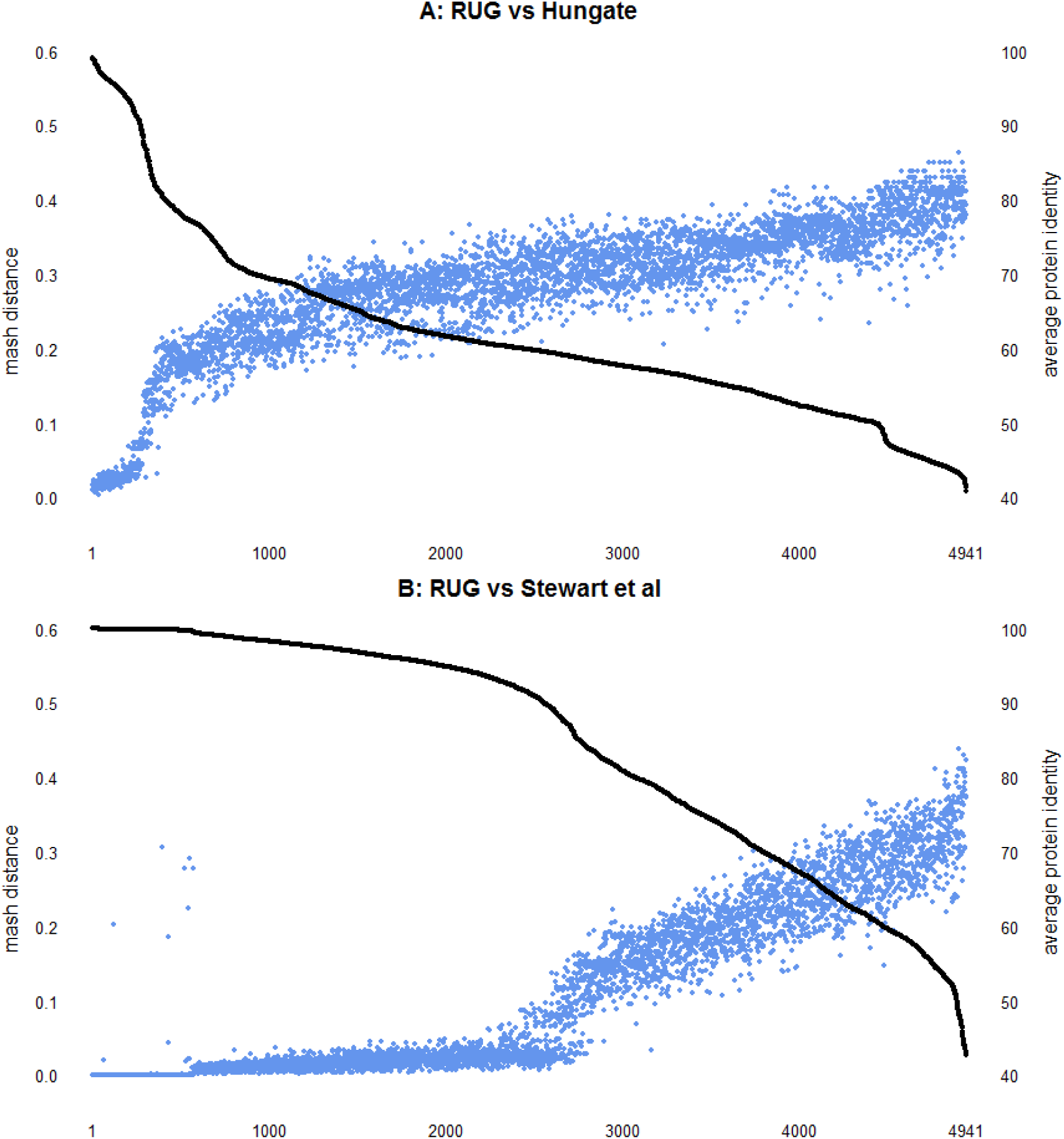
A comparison of the 4941 RUG dataset with the Hungate collection (A) and our previously published data from Stewart *et al* (B). Black line is average percentage protein identity with closest match (right-hand y-axis), and blue dots are mash distance (k=100,000) between RUG and the closest match (a measure of dissimilarity between two DNA sequences). As expected, a high protein identity relates to a low mash distance, and vice versa. The RUGs are sorted *interpedently* for figures A and B, by average protein identity.

### Assembly of long-read sequence data from the cow rumen

In addition to the Illumina data (above), a single sample was also sequenced on the Oxford Nanopore MinION sequencer. Three flowcells produced 11.4Gb of data with a read N50 of 11,585bp. The mean read length was 6144bp, which is short compared to other published projects^26^,^38^. We believe this is due to short DNA fragments and nicks in the DNA created by the bead-beating step of the DNA extraction.

We assembled the reads using Canu^39^, producing 1923 contigs greater than 1kb. The resulting assembly is 178Mb in length with an N50 of 268kb. 50 of the assembled contigs are >= 500kb in length, 26 are >= 1Mb in length, and five are >= 3Mb in length. Regardless of length, Canu predicted 31 of the contigs to be circular. A comparison of the Nanopore assembly statistics with those of 282 Illumina assemblies can be seen in Figure 6. The Nanopore assembly was created with a minimum contig length of 1kb, therefore the Illumina assemblies were similarly limited prior to analysis.

**Figure 6.**
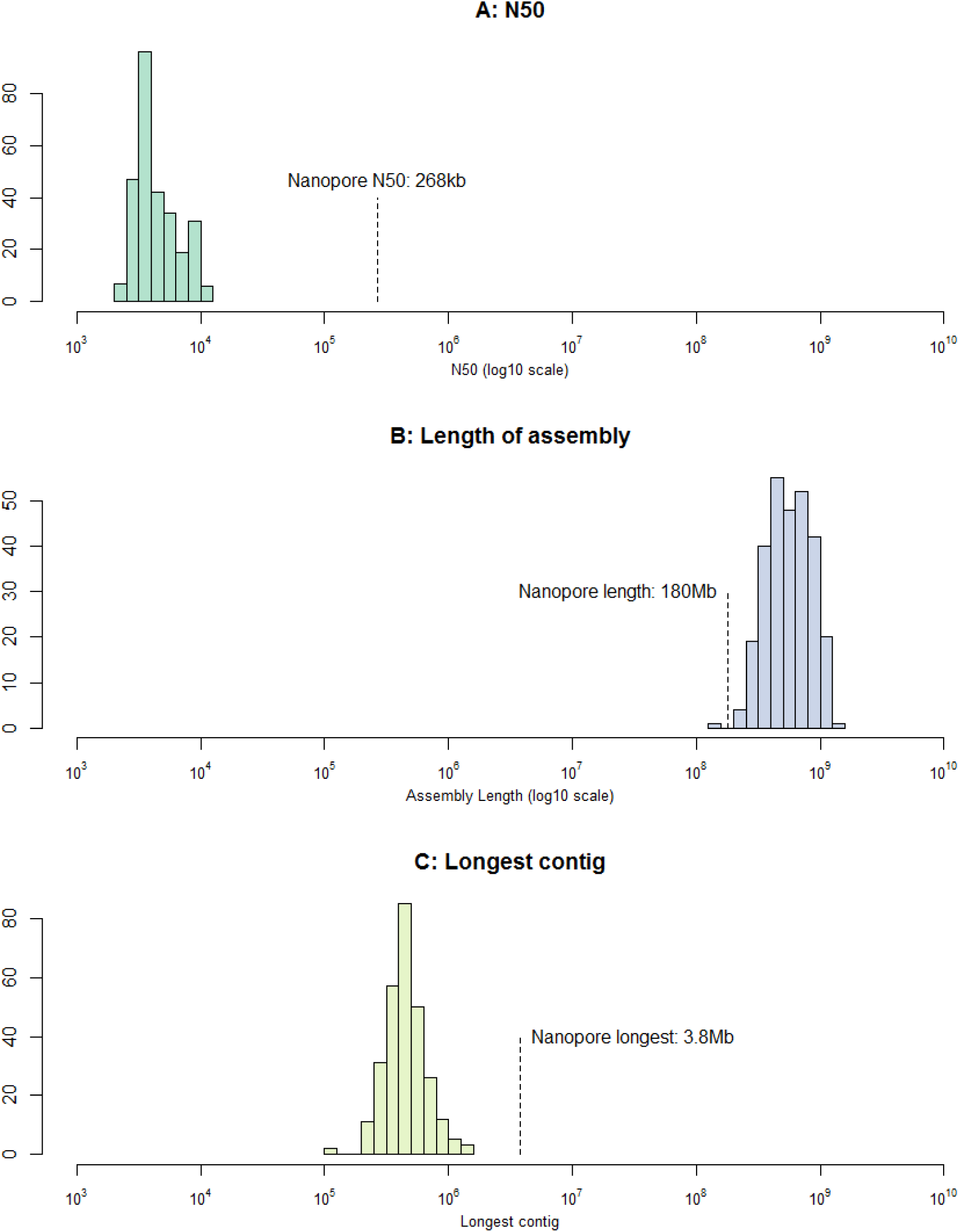
A comparison of Illumina and Nanopore metagenomic assembly statistics. The coloured histograms show the distribution of statistics for 282 Illumina assemblies, and the single Nanopore assembly is highlighted. **A)** N50; **B)** total length of the assembly; and **C)** length of the longest contig

As can be seen, the Nanopore assembly N50 of 268kb is over 56-times longer than the average Illumina assembly (4.7kb); whilst the Illumina assemblies are often longer (average 600Mb), the Nanopore assembly (at 178Mb in length) is not the shortest of the assemblies we produced; and the Nanopore assembly produced the longest contig at 3.8Mb, seven-times longer than the average for the Illumina assemblies (479kb) and 2.74-times longer than the longest single Illumina contig (1.38Mb – one of thirteen contigs from the 99.19% complete “uncultured *Bacteroidia* bacterium RUG14538”).

These results demonstrates that long read data are able to produce longer, more continuous metagenomic assemblies compared to short-read Illumina data, with only a minor reduction in the total assembly length.

A major issue with single-molecule sequencing technologies is the presence of post-assembly insertions and deletions (indels)^40^. Whilst Canu offers read-correction as part of the assembly process, this is often not enough to remove all indels. Detecting sequencing errors without a ground truth dataset is difficult; however, we hypothesized that in microbial genomes, the majority of indels would create premature stop-codons in open-reading frames (ORFs) and therefore gene prediction tools (we used Prodigal^41^) would produce truncated proteins. Examining the ratio between predicted protein length and their top-hit from UniProt should reveal if there are problems with indels – a dataset with no indels would produce a tight distribution around 1. We applied this to the Nanopore assembly before and after several correction algorithms, and include data from the Illumina RUGs as a control. The results are in Figure 7.

**Figure 7.**
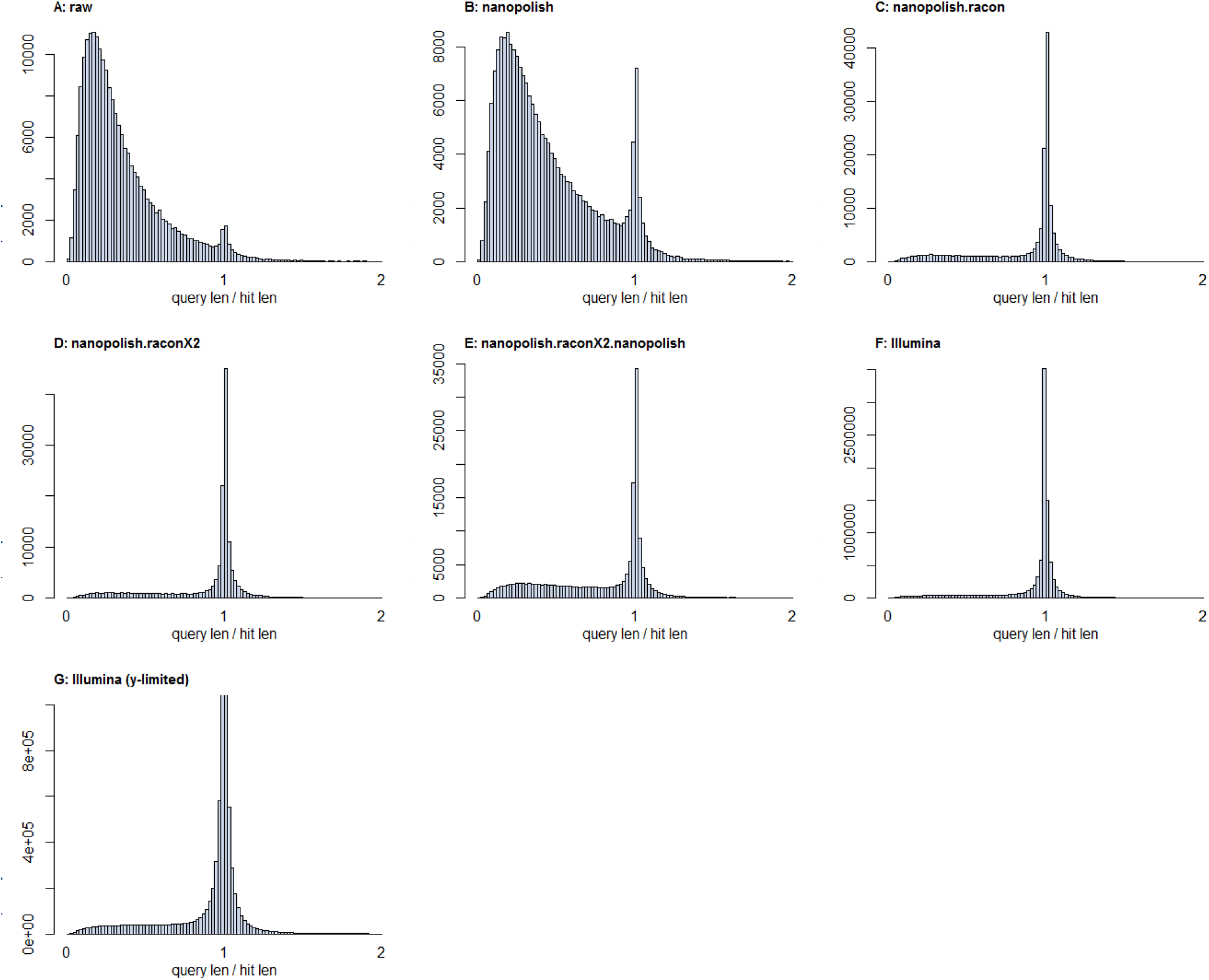
Histograms of predicted protein length vs length of the top hit in UniProt. Perfect predictions should show a tight distribution around 1. **A)** the raw assembly from Canu; **B)** after one round of Nanopolish; **C)** after one round of Nanopolish and one round of Racon; **D)** after one round of Nanopolish and two rounds of Racon; **E)** after one round of Nanopolish, two rounds of Racon, and a further round of Nanopolish; **F)** data from the 4941 RUGs; **G)** data from the 4941 RUGs with a limited y-axis, to highlight the long tail of short proteins remaining.

Clearly the raw assembly from Canu contains significant numbers of errors that truncate protein predictions – this is particularly apparent when compared to the Illumina data. However, the first round of Nanopolish^42^ produces a notable improvement, and the first round of Racon^43^ (with Illumina data) produces a drastic improvement. A second round of Racon (with Illumina data) produces a very slight improvement, and a final round of Nanopolish makes things slightly worse. On this basis, we decided to work with contigs polished with one round of Nanopolish and two rounds of Racon.

Statistics for all contigs >= 500kb and all contigs predicted to be circular can be found in Supplementary data 4. The Nanopore assembly contains several single contigs that we predict are complete or near-complete circular whole chromosomes.

*Prevotella copri* nRUG14950 (tig00000032) is a single contig of 3.8Mb which most closely resembles *Prevotella copri* DSM 18205, and which shows high similarity to RUG14032. *Prevotella copri* nRUG14950 is predicted to be 98.48% complete by CheckM^44^, with a contamination score of 2.03%; whereas RUG14032 is estimated to be 96.62% complete and 1.35% contaminated. Comparative alignments between *Prevotella copri* nRUG14950, RUG14032 and *Prevotella copri* DSM 18205 can be seen in Figure 8. There is a clear and striking relationship between *Prevotella copri* nRUG14950 and RUG14032. These two genomes, both estimated to be near-complete, were assembled from different samples using different techniques, and sequenced with different sequencing technologies. That such different approaches produce such similar results supports the integrity and quality of these assemblies. Alignments between *Prevotella copri* nRUG14950 and *Prevotella copri* DSM 18205 show a number of differences between the newly assembled genome and that of the current reference assembly – overall the two genomes are highly similar, but with some rearrangements which may be attributed to strain differences.

**Figure 8.**
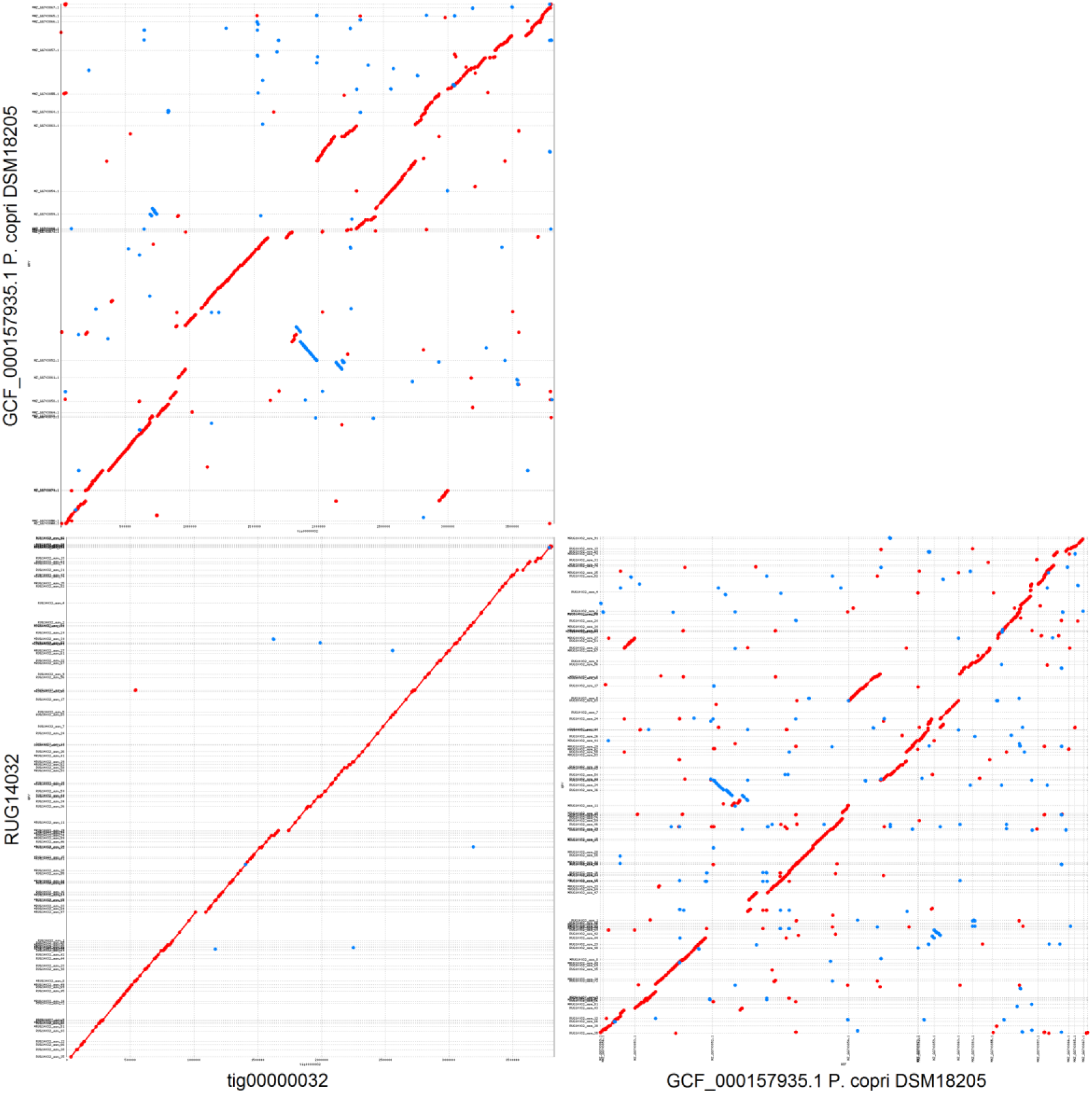
Whole genome alignments between *Prevotella copri* nRUG14950 (tig00000032), RUG14032 and *Prevotella copri* DSM18205. RUG14032 and *Prevotella copri* DSM18205 exist as unordered contigs whereas *Prevotella copri* nRUG14950 (tig00000032) is a single contig assembled from Nanopore data.

*Prevotella copri* nRUG14950 appears to capture as much of the genome as the reference assembly for *Prevotella copri* DSM 18205. At time of publication, 19 assemblies for *Prevotella copri* exist, with between 27 and 2159 scaffolds each. Our assembly of *Prevotella copri* nRUG14950, with only one contig and estimated to be highly complete, therefore represents the most continuous assembly of *Prevotella copri* to date, despite having been assembled from a metagenome.

Whilst *Prevotella copri* nRUG14950 has a high average protein identity with UniProt (91.42%), our Nanopore assembly contains other complete, predicted-circular assemblies with more distant relationships to public data. *Selenomonas spp.* nRUG14951 is a single contig of 3.1Mb in length, predicted to be circular, and with completeness and contamination statistics of 98.13% and 0.16% respectively. The most similar RUG is RUG10160, sharing a mean of 94% protein identity. RUG10160 is estimated 97.66% complete and 0% contaminated. However, the closest public reference genome is *Selenomonas ruminantium* GACV-9, part of the Hungate collection, which shares only ∼64% protein identity with *Selenomonas spp.* nRUG14951. There exists a good whole-genome alignment between *Selenomonas spp.* nRUG14951 and RUG10160 (Figure 9), albeit with some evidence of re-arrangements, and some small sections of the genome that are only captured by the Nanopore assembly. Whole genome alignments between nRUG14951, RUG10160 and *Selenomonas ruminantium* GACV-9 (Figure 9) suggest that the genomes share a similar genomic organisation, though these are weaker than for *Prevotella copri* (above), as would be expected between more distantly related organisms. A similar story emerges for *Lachnospiraceae bacterium* nRUG14952, another long (2.5Mb), circular, near-complete genome (95.46%) with a highly similar RUG (RUG13141, 96% protein identity) and a more distantly related public reference genome (*Lachnospiraceae bacterium* KHCPX20, 63% protein identity). Here, however, there are more pronounced differences between *Lachnospiraceae bacterium* nRUG14952 and RUG13141, with the Nanopore assembly containing larger regions of the genome that are not present in RUG13141 (Figure 10). The data suggest that both *Selenomonas spp.* nRUG14951 and *Lachnospiraceae bacterium* nRUG14952 represent entire bacterial genomes assembled as single contigs, and both now represent the best quality assembly for these species.

**Figure 9.**
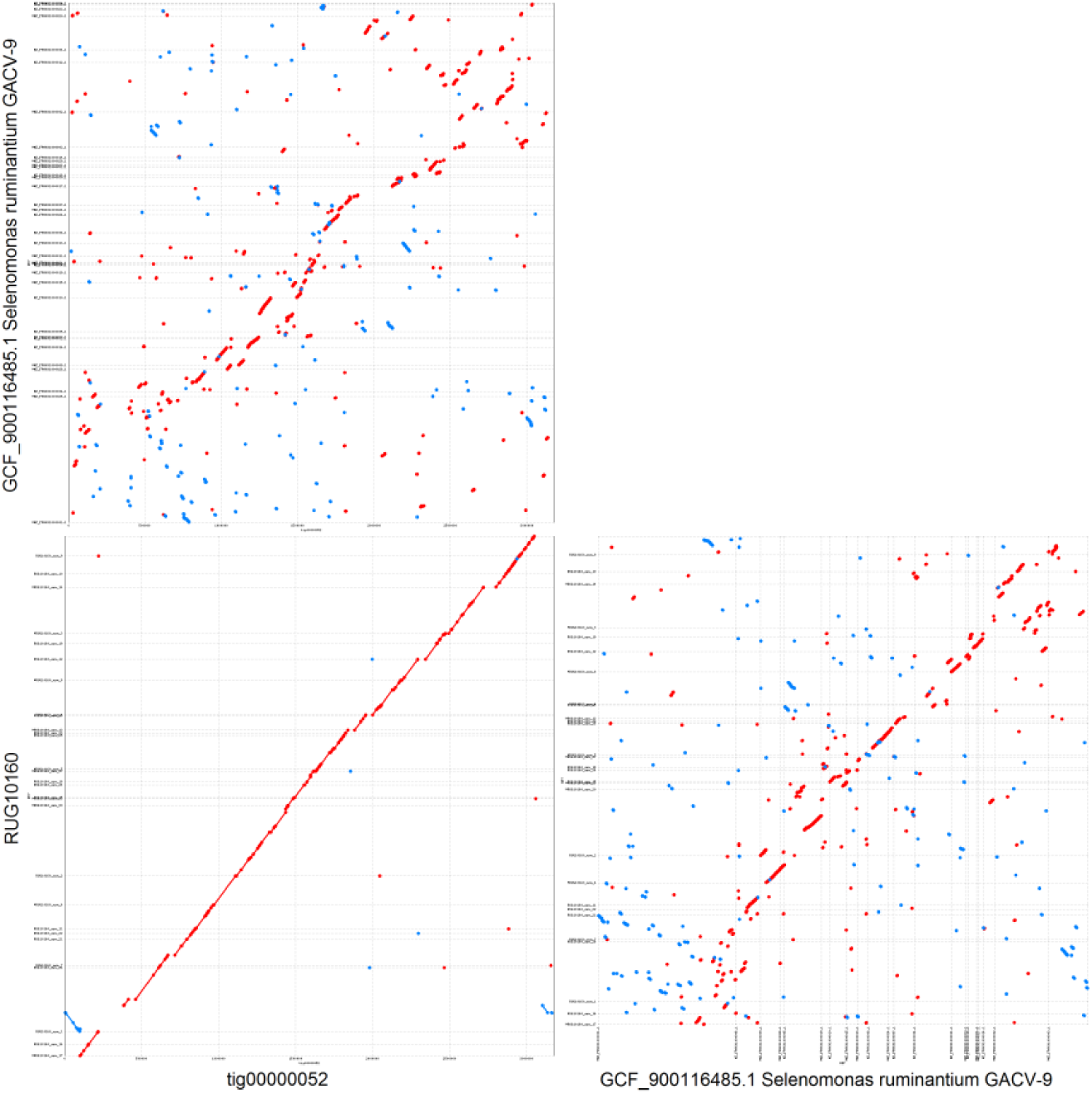
Whole genome alignments between *Selenomonas spp.* nRUG14951 (tig00000052), RUG110160 and *Selenomonas ruminantium* GACV-9. RUG110160 and *Selenomonas ruminantium* GACV-9 exist as unordered contigs whereas *Selenomonas spp* nRUG14951 (tig00000052) is a single contig assembled from Nanopore data.

**Figure 10.**
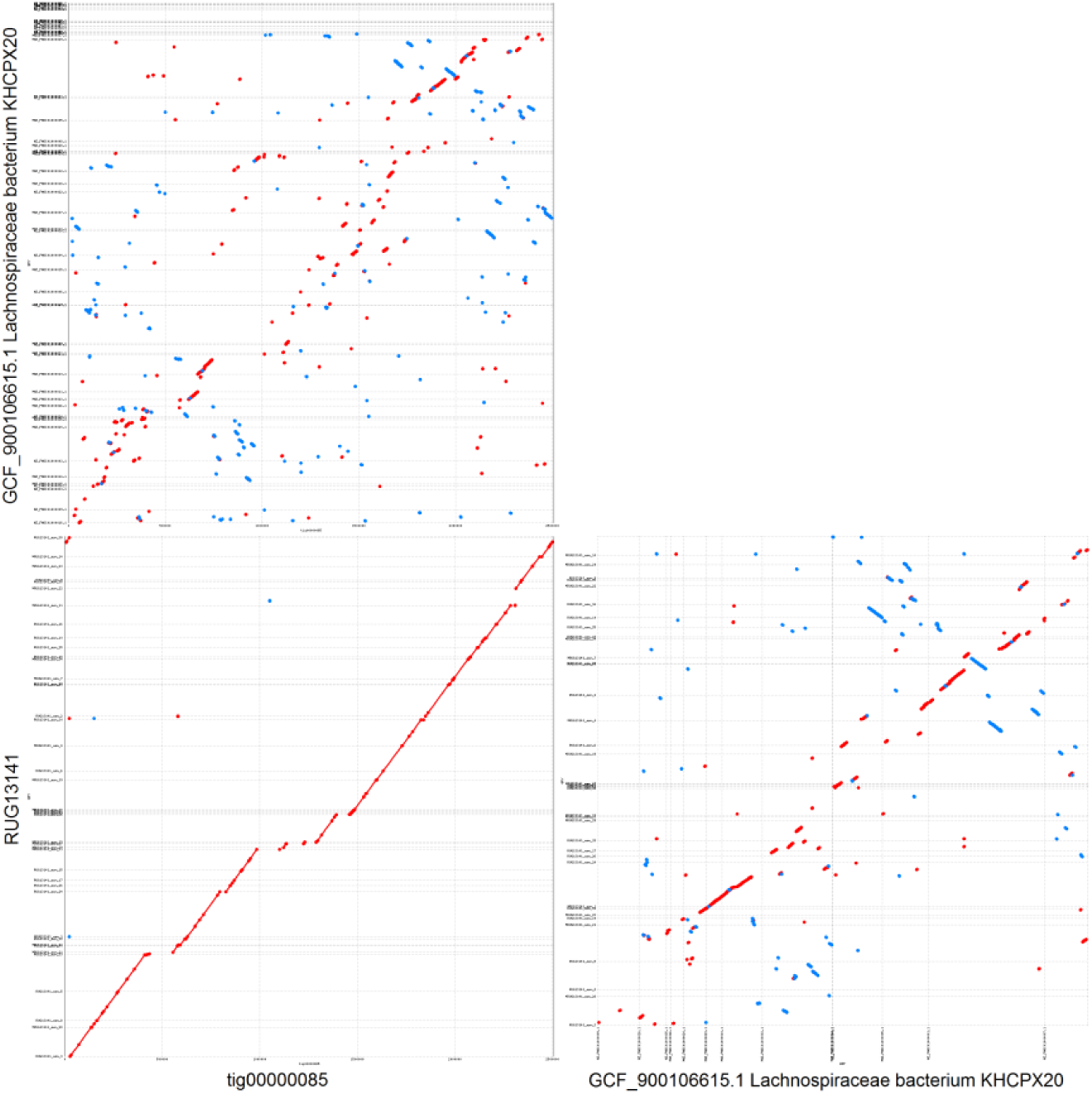
Whole genome alignments between *Lachnospiraceae bacterium* nRUG14952 (tig00000085), RUG13141 and *Lachnospiraceae bacterium* KHCPX20. RUG13141 and *Lachnospiraceae bacterium* KHCPX20 exist as unordered contigs whereas *Lachnospiraceae bacterium* nRUG14952 (tig00000085) is a single contig assembled from Nanopore data.

The above analysis demonstrates that our Nanopore assembly contains at least three near-complete, single-contig assemblies of novel rumen bacteria. Additionally, the remainder of the assembly contains highly continuous contigs that represent large portions of novel bacterial chromosomes. Some contigs may represent whole chromosomes, but not whole genomes, as some species of *Prevotella* have been shown to have two chromosomes^45^. These results taken together demonstrate the power of long reads for assembling complete, whole chromosomes from complex metagenomes.

### A dataset of rumen proteins for functional and proteomic analysis

We have created a non-redundant dataset of rumen proteins from the 4941 RUGs and 460 publicly available Hungate collection genomes, following the model of UniRef^46^ – clustering the protein set at 100%, 90% and 50% identity. We refer to these datasets as RumiRef100, RumiRef90 and RumiRef50.

The 4941 RUGs contain 10.69 million predicted proteins and the 460 Hungate collection genomes contain a further 1.29 million, resulting in a redundant dataset of 11.98 million proteins. This collapses to 9.45 million unique protein sequences at 100% identity, 5.69 million clusters at 90% identity and 2.45 million clusters at 50% identity.

Focusing on RumiRef90, 4.1 million of the 5.69 million clusters are singletons, 3.53 million from the RUG dataset and 566k from the Hungate genomes. Of the 1.58 million clusters with more than one member, 1.33 million contain only proteins from the RUGs, whereas 257K clusters contain at least one Hungate collection protein. Of those, 107 thousand clusters contain only Hungate collection proteins, and 149 thousand clusters contain a mix of RUG and Hungate proteins.

All 10.69 million predicted proteins from the RUGs have been compared to KEGG^47^, 460 public genomes from the Hungate collection, UniRef100, UniRef90 and UniRef50. The mean protein identity of the top hit for these databases is 55.88%, 63.58%, 67.52%, 67.25% and 59.97% respectively. These data provide the largest, most comprehensive and richly annotated protein dataset from the rumen to date. Together, these results indicate that the RUGs contain a huge amount of novelty at the protein level compared to the Hungate collection and other publicly available rumen microbial genomes, and both the redundant and non-redundant datasets presented here will form an important reference dataset for future functional and proteomic studies in the rumen.

The RUG proteins were compared to the CAZy^48^ database of 31^st^ July 2018 using dbCAN2^49^. Of the 10.69 million proteins, 442917 are predicted to be involved in carbohydrate metabolism, including 235001 glycoside hydrolases, 120494 glycosyltransferases, 55523 carbohydrate esterases, 23928 proteins with carbohydrate binding modules, 6834 polysaccharide lyases, 907 proteins with predicted auxiliary activities, 80 proteins with a predicted cohesin domain, and 150 proteins with an S-layer homology module (SLH). Cohesin and SLH domains are associated with cellulosomes, enzyme complexes produced by cellulolytic microorganisms to digest cellulose^50^.

The novelty of the predicted CAZymes relative to the current CAZy database can be seen in Figure 11. None of the eight classes of carbohydrate active enzymes displays an average protein identity greater than 60% indicating that CAZy poorly represents the diversity of CAZymes encoded in the genomes of ruminant microbes.

**Figure 11.**
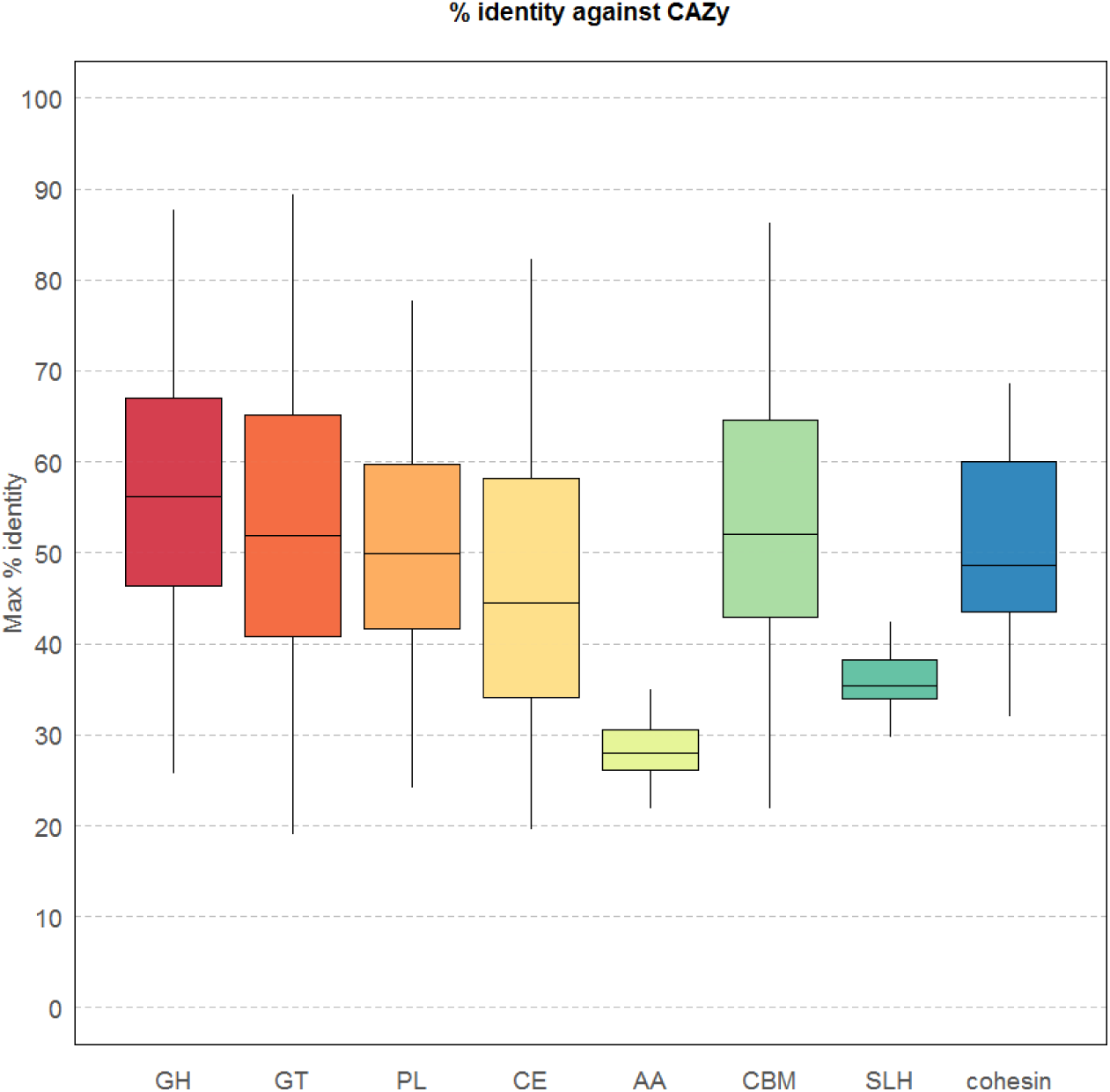
Maximum percentage identity between CAZyme-predicted proteins from the RUGs and the CAZy database. GH=glycoside hydrolase; GT=glycosyl transferase; PL=polysaccharide lyase; CE=carbohydrate esterase; AA=auxiliary activities; CBM=carbohydrate binding module; SLH=S-layer homology domain; cohesin=cohesin domain

Of particular note is the class AA “auxiliary activities”, with an average protein identity of less than 30% between CAZy and the RUG CAzymes. AA was created by CAZy to classify ligninolytic enzymes and lytic polysaccharide mono-oxygenases (LPMOs). Ligninolytic enzymes are involved in the breakdown of lignin, an organic structural polymer present in plant cell walls and wood; LPMOs are a class of oxidative enzymes, involved in the breakdown of cellulose by activation of oxygen. The rumen is a largely anaerobic environment, though some oxygen is present^51^, and this may explain the low similarity between rumen LPMOs and those publicly available. The SLH-encoding proteins also show a low level of protein similarity, indicating that AA and SLH class CAZymes from the rumen are particularly poorly represented by CAZy.

The distribution of CAZymes across 12 different phyla and the group of “unknown” bacteria can be seen in Figure 12. The *Bacteroidetes* (3.9 million) and *Firmicutes* (5.3 million) together contribute the largest number of proteins to our dataset; however, whereas 5.7% of the proteome of *Bacteroidetes* is devoted to CAZyme activity, in *Firmicutes* the figure is 3.2%. *Fibrobacteres* devote the highest percentage of their proteome to carbohydrate metabolism (over 6.6%), as is expected due to their fibre-attached, high cellulolytic activity. Only a few studies exist on the role of *Planctomycetes* in the rumen^52–54^, however whilst they contribute a relatively low number of proteins in our dataset (30172), just over 5% of those proteins are predicted to be CAZymes, suggesting a role in and adaptation to carbohydrate metabolism. Most phyla encode relatively similar proportions of each of the eight classes of CAZyme; however, the archaea encode proportionally more glycosyl transferases, and the *Fibrobacteres* encode proportionally more carbohydrate binding molecules. The *Tenericutes* encode only glycosyl transferases, though these make up only 0.3% of their proteome. 79 out of 80 cohesin-containing proteins are encoded by the *Firmicutes* (the remaning one is encoded by an unknown bacterium), as are 101 out of 149 SLH-domain containing proteins. Both are components of cellulosomes, abundant within the *Clostridiales.* Finally, 329910 proteins are encoded by unknown bacteria, and 5% of these are predicted to be CAZymes, suggesting that many bacteria with important roles in carbohydrate metabolism remain to be identified in the rumen.

**Figure 12.**
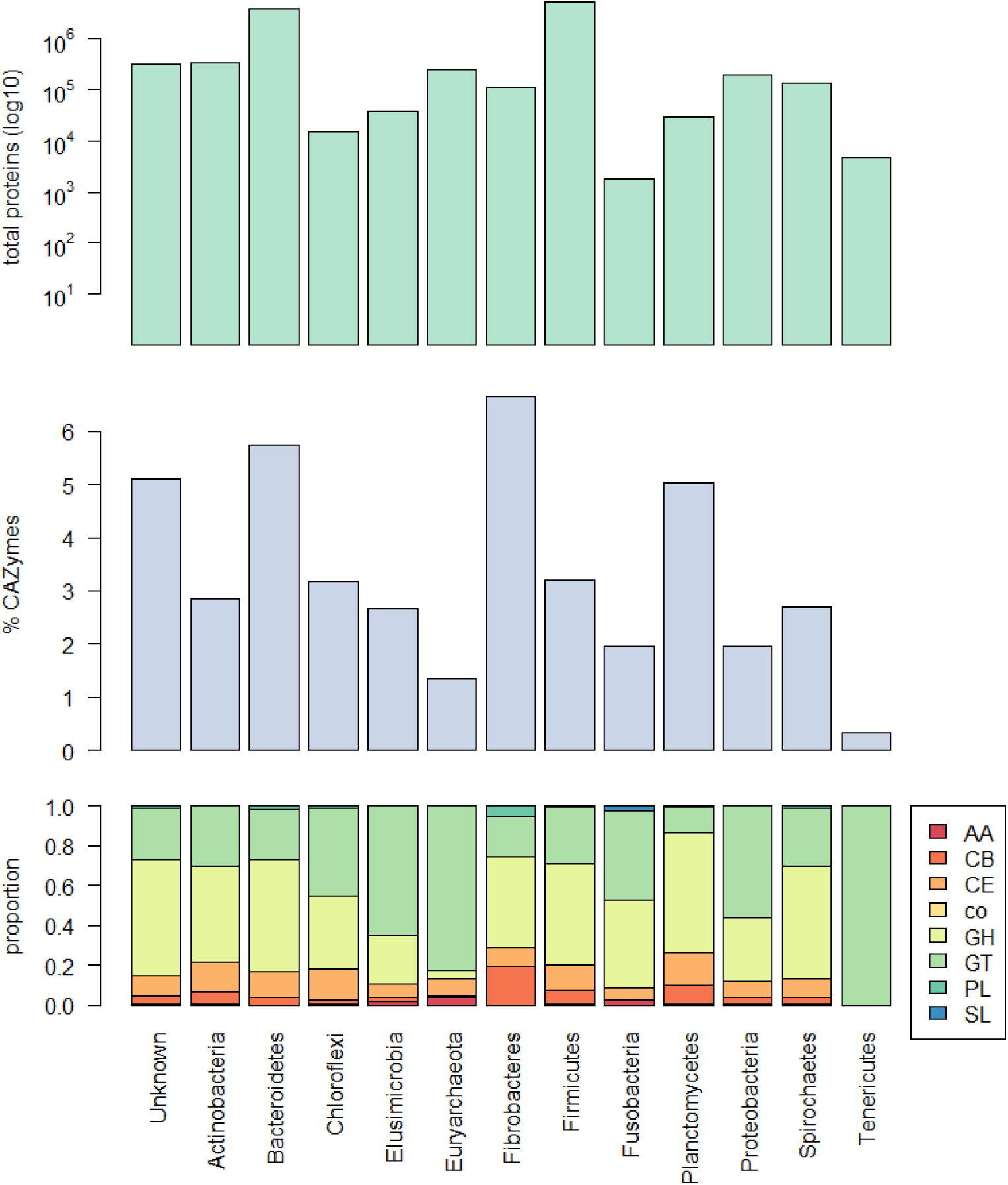
Top: Total number of proteins for 12 phyla and the group of unknown bacteria; Middle: percentage of the proteome predicted to be CAZymes; Bottom: distribution of eight CAZyme classes as a proportion of the total number of predicted CAZymes. GH=glycoside hydrolase; GT=glycosyl transferase; PL=polysaccharide lyase; CE=carbohydrate esterase; AA=auxiliary activities; CB=carbohydrate binding module; SL=S-layer homology domain; co=cohesin domain

Polysaccharide utilisation loci (PUL) are loci of co-located genes involved in the binding and degradation of specific carbohydrates^55^, and are characterised in *Bacteroidetes* by a susC-susD gene pair with predicted CAZymes encoded nearby. There are 1707 *Bacteroidetes* genomes in the RUGs, and additionally we have a whole genome of *Prevotella copri* from the Nanopore assembly. These 1708 genomes were subject to prediction of polysaccharide utilisation loci (PUL) using our pipeline PULpy^56^.

Of the 1708 genomes, 1469 are predicted to have at least one PUL, and in total there are 15,629 separate loci involving 88260 proteins. The highest number of PUL per genome are 52 PUL for RUG13980 and 50 for RUG10279, both labelled uncultured *Prevotellaceae;* both of these genomes are closely related to *Prevotella multisaccharivorax*, known to be able to utilise multiple carbohydrate substrates^57^. *Prevotella copri* nRUG14950 encodes 25 PUL, including one that spans over 50kb – predicting such long loci is only possible in highly continuous genome assemblies.

The current version of PULDB^58^, the public database of PUL, contains 30,232 PUL; and recently we released data on over 96,000 PUL from all public *Bacteroidetes* genomes^56^. The additional 15,629 PUL predicted here ensures that ruminant-microbe-encoded PUL are represented in the public domain, and these loci will be a hugely interesting resource for researchers interested in carbohydrate utilisation in the rumen.

## Discussion

Arguably the rumen is one of the most economically important microbiomes on the planet, converting low value cattle feed into high value milk and meat protein that is consumed by billions of people worldwide; as part of this process, methane is released by the rumen methanogenic archaea, significantly contributing to greenhouse gas emissions and climate change. Understanding and studying the structure and function of the rumen microbiome is of clear importance.

Recent studies have released large amounts of genomic and metagenomic data derived from the ruminant microbiome, including over 1300 draft and complete genomes. Here we present almost 5000 additional genomes, from both Illumina and Nanopore sequencing data, and demonstrate that there remain thousands of novel strains of bacteria and archaea, and millions of novel proteins, in the rumen. The Hungate collection is a collection of 410 bacteria and archaea in culture, yet we have shown that these genomes represent only a fraction of the diversity present in our samples. It is absolutely vital that more of these strains are brought into culture, so we can study their function *in vitro* and *in vivo* and gain mechanistic insight into the structure and function of the rumen microbiome. In particular, if we are to design rational interventions to manipulate rumen feed-conversion or methane emissions, we will need to understand which microbes are present, which substrates they feed on, and how they interact with one another and the host. For these questions we not only need to understand their genomic organisation, but how they behave and interact. Sequencing and assembling their genomes, as we have done in this study, is the first step towards improved culture collections and future manipulation of the rumen microbiome for human benefit.

The activity of the rumen microbiota is largely defined by the activity of proteins encoded in their genomes, and as researchers produce more proteomic data, it is vital that good protein reference datasets are available. We have demonstrated that public datasets and databases poorly represent the protein diversity of the rumen. Here we present the largest redundant and non-redundant rumen microbial protein prediction datasets to date, and provide rich annotation against public protein, pathway and enzyme databases. This rich annotation allows researchers to predict the function of each protein, and better assess the functional consequences of changes in the rumen proteome.

We have used two different sequencing technologies to assemble microbial genomes. The benefits of Illumina data are that it is both cheap to produce large amounts of sequence data, and that sequence data is highly accurate. Advances in bioinformatics allow us to assemble and bin these data into high quality putative genomes; however, the assemblies of these genomes are still highly fragmented. Nevertheless, they are very useful for assessing the functional capacity of microbial species and for discovering new strains and species of microbe. In contrast, Nanopore sequencing data is comparatively more expensive, yet the longer reads allow more complete genome fragments to be assembled, and we demonstrate in this paper that whole bacterial chromosomes can be assembled into single contigs. Despite the raw error rate and predominance of indels, these can largely be corrected using bioinformatics techniques and additional Illumina data. As long reads become cheaper and quality improves, we expect them to become the method of choice for sequencing metagenomes, with the added benefit that more metagenome-assembled genomes will be highly continuous, single-contig assemblies.

## Methods

### Metagenomic samples

Animal experiments were conducted at the Beef and Sheep Research Centre of Scotland’s Rural College (SRUC). The experiment was approved by the Animal Experiment Committee of SRUC and was conducted in accordance with the requirements of the UK Animals (Scientific Procedures) Act 1986.

The data were obtained from three cross breeds: Aberdeen Angus, Limousin and Charolais and one pure breed: Luing. As previously described, the animals were slaughtered in a commercial abattoir where two post-mortem digesta samples (approximately 50 mL) were taken immediately after the rumen was opened to be drained^59^,^60^. DNA extraction was carried out following the protocol of Yu and Morrison^61^ and based on repeated bead beating plus column filtration. Illumina TruSeq libraries were prepared from genomic DNA and sequenced on an Illumina HiSeq 4000 by Edinburgh Genomics. For the MinION Nanopore sequencing, additional clean-up was carried out using phenol-chloroform extraction and a 1D library was prepared following Oxford Nanopore’s SQK-LSK108 1D ligation protocol. Three sequencing runs were carried out using FLOMIN-106 flow cells on a MinION MK1b in the Watson lab.

### Bioinformatics – metagenomic assembly and binning

In total, 282 samples were sequenced generating between 24 and 140 million 150bp paired-end reads per sample. The samples were sequenced in five batches of 48 samples and one batch of 42 samples (this 42-sample batch was the sole basis of Stewart *et al*).

Each sample was assembled and binned individually using coverage and content as previously described^11^. Briefly, each sample was assembled using idba_ud^15^ with the options --num_threads 16 --pre_correction -- min_contig 300. BWA MEM^62^ was used to map reads back to the filtered assembly and Samtools^63^ was used to convert to BAM format. Script jgi_summarize_bam_contig_depths from the MetaBAT2^18^ package was used to calculate coverage from the resulting BAM files. A co-assembly was also produced for each of the 6 batches of samples using MEGAHIT^16^ with options --kmin-1pass, -m 60e +10, --k-list 27,37,47,57,67,77,87, -- min-contig-len 1000, -t 16.

Metagenomic binning was applied to both single-sample assemblies and the coassemblies using MetaBAT2^18^, with options --minContigLength 2000, --minContigDepth 2. Single-sample binning produced a total of 37153 bins, and co-assembly binning produced a further 23335. All 60743 bins were aggregated and then dereplicated using dRep^64^. The dRep dereplication workflow was used with options dereplicate_wf -p 16 - comp 80 -con 10 -str 100 –strW 0. Thus, in pre-filtering, only bins assessed by CheckM as having both completeness ≥ 80%, and contamination ≤ 10% were retained for pairwise dereplication comparison. Bin scores were given as completeness - 5*contamination + 0.5*log(N50), and only the highest scoring RUG from each secondary cluster was retained in the dereplicated set. For our dataset, 4941 dereplicated RUGs were obtained, 4109 from the single-sample assemblies and 800 from the co-assembly.

Note that we operate a continuous de-replication workflow. Therefore all 913 of the RUGs (both MetaBAT2 and Hi-C) we previously published have been merged with the newer RUGs and de-replicated. Therefore, whilst some of the previously published RUGs exist in the newer dataset published here, many have been replaced by newer RUGs of higher quality.

The RUGs were compared to the Hungate collection and our previous dataset using DIAMOND and MASH^65^.

### Bioinformatics – assembly of Nanopore sequence data

The Nanopore reads were extracted and QC-ed using poRe^66^,^67^, and assembled using Canu^39^ with default settings and genomeSize=150Mb. The resulting assembly was analysed using MAGpy^19^. Whole-genome alignments were calculated using MUMmer^68^. The raw assembly was corrected using both Nanopolish^27^ and Racon^43^ (using Illumina data aligned to the Nanopore assembly with Minimap2 using short read settings (-x sr). Query vs subject length data were extracted from DIAMOND^31^ results and plotted using R.

### Bioinformatics – metagenomic assignment

The output of metagenomic binning is simply a set of DNA FASTA files containing putative genomes. These were all assessed for completeness and contamination using CheckM^44^. The 4941 best bins were analysed using MAGpy^19^, a Snakemake^69^ pipeline that runs a series of analyses on the bins, including: CheckM; prodigal^41^ protein prediction; Pfam_Scan^70^; DIMAOND^31^ search against UniProt TrEMBL; and Sourmash search against all public bacterial genomes. The MAGpy results were used to produce a putative taxonomic assignment for each bin. A phylogenetic tree consisting of the RUGs and 460 genomes from the Hungate 1000 collection, produced using PhyloPhlAn^32^, was visually inspected using FigTree, iTol^71^ and GraPhlAn^72^. Annotations were improved where they could be - for example where MAGpy had only assigned a taxonomy at the Family level, but where a genome clustered closely with a Hungate 1000 genome annotated at the species level, the annotation was updated. The tree was also re-rooted manually at the Bacteria/Archaea branch using FigTree.

### Bioinformatics – proteome analysis

Proteins were predicted using Prodigal with option –p meta. Using DIAMOND, each protein was searched against KEGG (downloaded on Sept 15^th^ 2018), UniRef100, UniRef90 and UniRef50 (downloaded Oct 3^rd^ 2018), and CAZy (dbCAN2 version, 31/07/2018). The protein predictions were clustered by CD-HIT^73^ at 100%, 90% and 50% identity, mirroring similar methods at UniRef.

All protein predictions were searched against the CAZy database using dbCAN2^49^ and HMMER^74^, and polysaccharide utilisation loci (PUL) were predicted for *Bacteroidetes* RUGs using PULpy^56^.

## Data Availability

Raw sequence reads are in the process of being deposited in ENA. All assemblies and protein predictions are available at DOI: 10.7488/ds/2470.

## Supporting information

## Acknowledgements

The Rowett Institute and SRUC are core funded by the Rural and Environment Science and Analytical Services Division (RESAS) of the Scottish Government. The Roslin Institute forms part of the Royal (Dick) School of Veterinary Studies, University of Edinburgh. This project was supported by the Biotechnology and Biological Sciences Research Council (BBSRC; BB/N016742/1, BB/N01720X/1), including institute strategic programme and national capability awards to The Roslin Institute (BBSRC: BB/P013759/1, BB/P013732/1, BB/J004235/1, BB/J004243/1); and by the Scottish Government as part of the 2016–2021 commission.

## Author contributions

MW, RR and AWW conceived of the study and supervised the project. RDS carried out all bioinformatics work on the Illumina data and AW carried out all bioinformatics work on the Nanopore data. MA carried out all laboratory work. All of the authors contributed ideas, co-wrote the paper, and reviewed and approved the manuscript.

## Competing interests

The authors declare no competing interests.

